# Deep reinforcement learning for modeling human locomotion control in neuromechanical simulation

**DOI:** 10.1101/2020.08.11.246801

**Authors:** Seungmoon Song, Łukasz Kidziński, Xue Bin Peng, Carmichael Ong, Jennifer Hicks, Sergey Levine, Christopher G. Atkeson, Scott L. Delp

## Abstract

Modeling human motor control and predicting how humans will move in novel environments is a grand scientific challenge. Despite advances in neuroscience techniques, it is still difficult to measure and interpret the activity of the millions of neurons involved in motor control. Thus, researchers in the fields of biomechanics and motor control have proposed and evaluated motor control models via neuromechanical simulations, which produce physically correct motions of a musculoskeletal model. Typically, researchers have developed control models that encode physiologically plausible motor control hypotheses and compared the resulting simulation behaviors to measurable human motion data. While such plausible control models were able to simulate and explain many basic locomotion behaviors (e.g. walking, running, and climbing stairs), modeling higher layer controls (e.g. processing environment cues, planning long-term motion strategies, and coordinating basic motor skills to navigate in dynamic and complex environments) remains a challenge. Recent advances in deep reinforcement learning lay a foundation for modeling these complex control processes and controlling a diverse repertoire of human movement; however, reinforcement learning has been rarely applied in neuromechanical simulation to model human control. In this paper, we review the current state of neuromechanical simulations, along with the fundamentals of reinforcement learning, as it applies to human locomotion. We also present a scientific competition and accompanying software platform, which we have organized to accelerate the use of reinforcement learning in neuromechanical simulations. This “Learn to Move” competition, which we have run annually since 2017 at the NeurIPS conference, has attracted over 1300 teams from around the world. Top teams adapted state-of-art deep reinforcement learning techniques to produce complex motions, such as quick turning and walk-to-stand transitions, that have not been demonstrated before in neuromechanical simulations without utilizing reference motion data. We close with a discussion of future opportunities at the intersection of human movement simulation and reinforcement learning and our plans to extend the Learn to Move competition to further facilitate interdisciplinary collaboration in modeling human motor control for biomechanics and rehabilitation research.

## Introduction

Predictive neuromechanical simulations can produce motions without directly using experimental motion data. If the produced motions reliably match how humans move in novel situations, predictive simulations can be used to accelerate research on assistive devices, rehabilitation treatments, and *Preprint submitted to Journal Name August 11, 2020* physical training. Neuromechanical models represent the neuro-musculo-skeletal dynamics of the human body and can be simulated based on physical laws to predict body motions (Fig. 1). Although advancements in musculoskeletal modeling [1, 2] and physics simulation engines [3, 4, 5] allow us to simulate and analyze observed human motions, understanding and modeling human motor control remains a hurdle for accurately predicting motions. In particular, it is difficult to measure and interpret the biological neural circuits that underlie human motor control. To overcome this challenge, one can propose control models based on key features observed in animals and humans and investigate these models in neuromechanical simulations by comparing the simulation results to human data. With such physiologically plausible neuromechanical control models, today we can simulate many aspects of human motions, such as locomotion, in a predictive manner [6, 7, 8]. Despite this progress, developing controllers for more complex tasks, such as adapting to dynamic environments and those that require long-term planning, remains a challenge.

**Figure 1:**
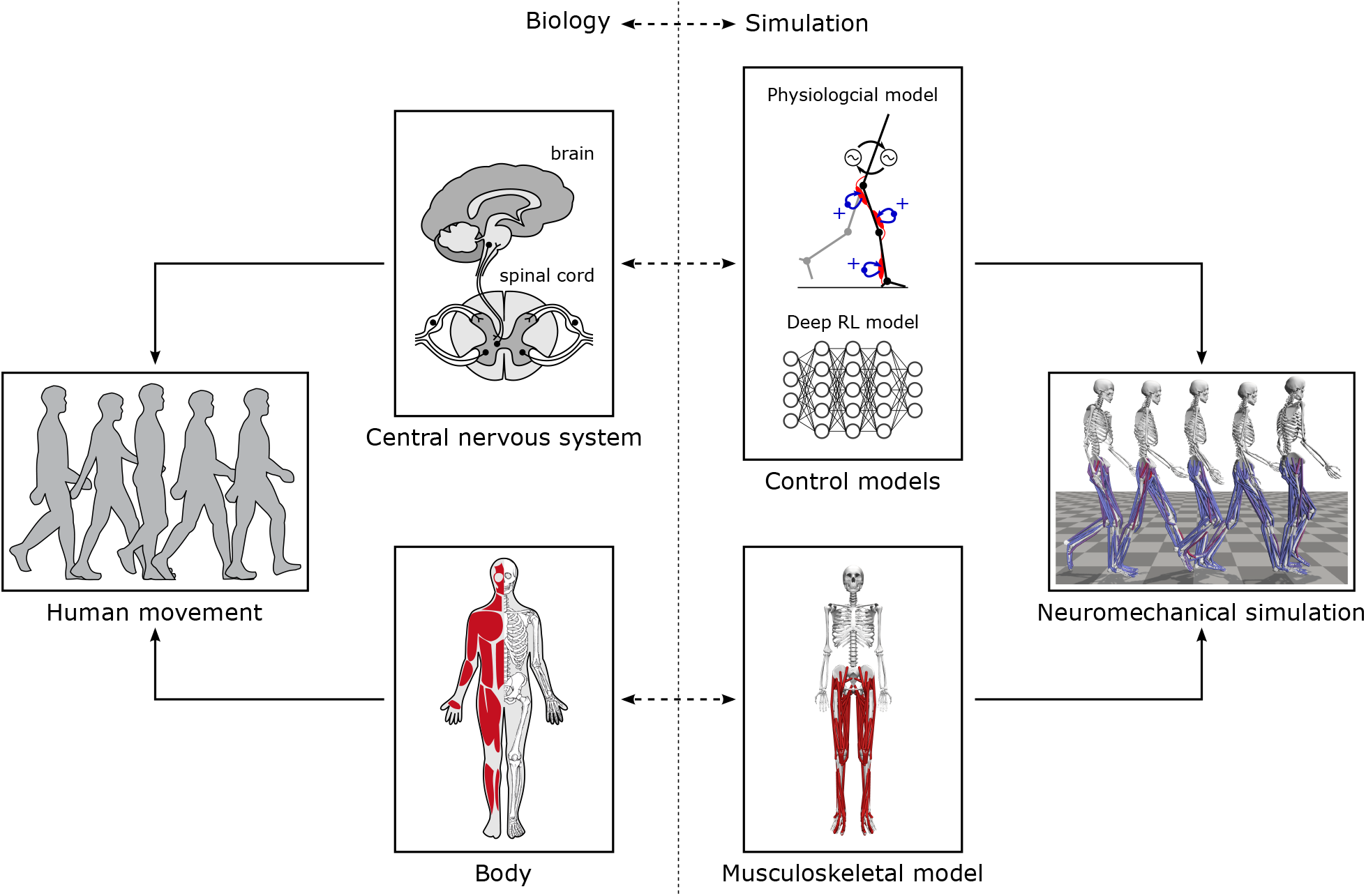
Neuromechanical simulation. A neuromechanical simulation consists of a control model and a musculoskeletal model that represent the central nervous system and the body, respectively. The control and musculoskeletal models are forward simulated based on physical laws to produce movements.

Training artificial neural networks using deep reinforcement learning (RL) in neuromechanical simulations provides an alternative way of developing control models. In contrast to developing a control model that captures certain physiological features and then running simulations to evaluate the results, deep RL can be thought of as training a black-box controller that produces motions of interest. Recent breakthroughs in deep learning make it possible to develop controllers with high-dimensional inputs and outputs that are applicable to human musculoskeletal models. Despite the discrepancy between artificial and biological neural networks, such means of developing versatile controllers could be useful in investigating human motor control [9]. For instance, one could train a controller (i. e., a policy in RL terminology) implemented on an artificial neural network using deep RL in a physiologically plausible simulation environment, and then investigate or reverse engineer the resulting network. One could also train controllers to mimic human motion (e.g., using imitation learning, where a controller is trained to replicate behaviors demonstrated by an expert [10]) or integrate an existing neuromechanical control model with artificial neural networks to study certain aspects of human motor control. While there are recent studies that used deep RL to produce human-like motions with musculoskeletal models [11, 12], little effort has been made to study the underlying control.

We organized the Learn to Move competition series to facilitate developing control models with advanced deep RL techniques in neuromechanical simulation. It has been an official competition at the NeurIPS conference for the past three years. We provided the neuromechanical simulation environment, OpenSim-RL, and participants developed locomotion controllers for a human musculoskeletal model. In the most recent competition, NeurIPS 2019: Learn to Move - Walk Around, the top teams adapted state-of-the-art deep RL techniques and successfully controlled a 3D human musculoskeletal model to follow target velocities by changing walking speed and direction as well as transitioning between walking and standing. Some of these locomotion behaviors were demonstrated in neuromechanical simulations for the first time without using reference motion data. While the solutions were not explicitly designed to model human learning or control, they provide means of developing control models that are capable of producing realistic complex motions.

This paper reviews neuromechanical simulations and deep RL, with a focus on the materials relevant to modeling the control of human locomotion. First, we review control models of human locomotion that have been studied in computer simulations and discuss how to evaluate their physiological plausibility. By reviewing human control models, we hope to inspire future studies using computational approaches, such as deep RL, to encode physiologically plausible features. We also introduce deep RL approaches for continuous control problems (the type of problem we must solve to predict human movement) and review their use in developing locomotion controllers. Then, we present the Learn to Move competition and discuss the successful approaches, simulation results, and their implications for locomotion research. We conclude by suggesting promising future directions for the field and outline our plan to extend the Learn to Move competition.

### Computer simulations of human locomotion

This section reviews computer simulation studies that propose control models of human locomotion. We first present the building blocks of musculoskeletal simulations and their use in studying human motion. We next review the biological control hypotheses and neuromechanical control models that embed those hypotheses. We also briefly cover studies in computer graphics that have developed locomotion controllers for human characters. We close by discussing the means of evaluating the plausibility of control models and the limitations of current approaches.

#### Musculoskeletal simulations

A musculoskeletal model typically represents a human body with rigid segments and muscle-tendon actuators [13, 14, 15] (Fig. 2-a). The skeletal system is often modeled by rigid segments connected by rotational joints. Hill-type muscle models [16] are commonly used to actuate the joints, capturing the dynamics of biological muscles, including both active and passive contractile elements [17, 18, 19, 20] (Fig. 2-b). Hill-type muscle models can be used with models of metabolic energy consumption [21, 22, 23] and muscle fatigue [24, 25, 26] to estimate these quantities in simulations. Musculoskeletal parameter values are determined for average humans based on measurements from a large number of people and cadavers [27, 28, 29, 30] and can be customized to match an individual’s height, weight, or CT and MRI scan data [31, 32]. OpenSim [1], which is the basis of the OpenSim-RL package [33] used in the Learn to Move competition, is an open-source software package broadly used in the biomechanics community (e.g., it has about 40,000 unique user downloads) to simulate musculoskeletal dynamics.

**Figure 2:**
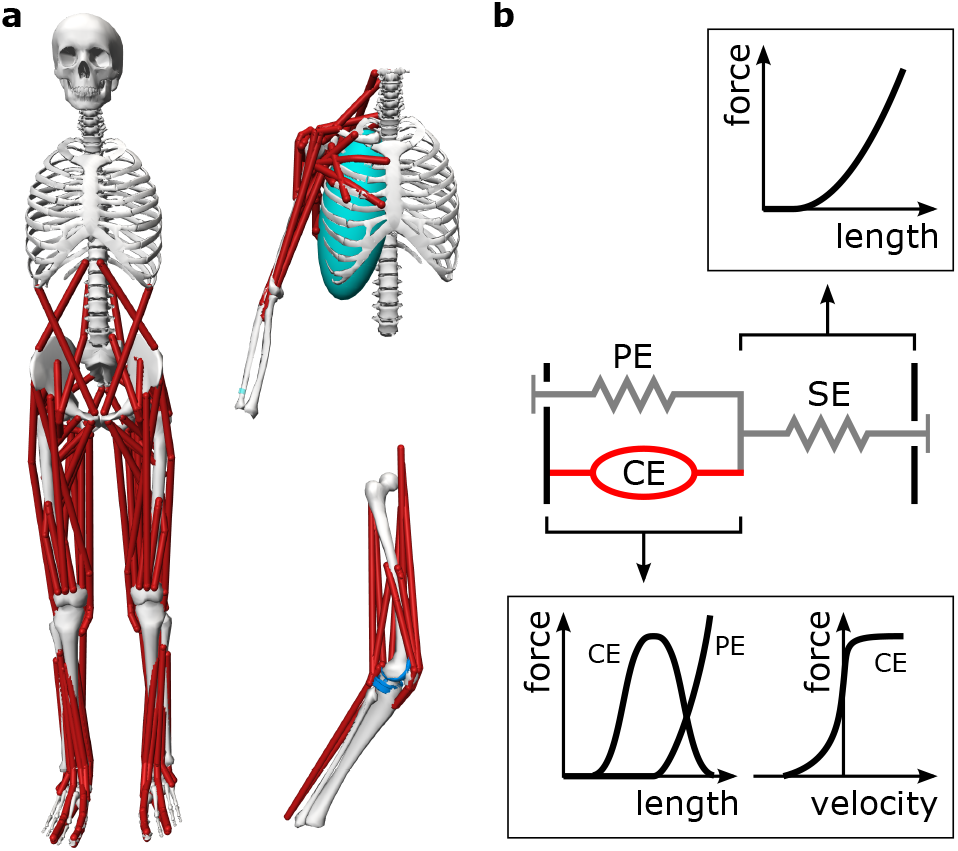
Musculoskeletal models for studying human movement. a. Models implemented in OpenSim [1] for a range of studies: lower-limb muscle activity in gait [34], shoulder muscle activity in upper-limb movements [15], and knee contact loads for various motions [35]. b. A Hill-type muscle model typically consists of a contractile element (CE), a parallel elastic element (PE), and a series elastic element (SE). The contractile element actively produces contractile forces that depend on its length and velocity and are proportional to the excitation signal. The passive elements act as non-linear springs where the force depends on their length.

Musculoskeletal simulations have been widely used to analyze recorded human motion. In one common approach, muscle activation patterns are found through various computational methods to enable a musculoskeletal model to track reference motion data, such as motion capture data and ground reaction forces [36, 37, 38], while achieving defined objectives, like minimizing muscle effort. The resulting simulation estimates body states, such as individual muscle forces, that are difficult to directly measure with an experiment. This approach has been extensively used to analyze human locomotion [36, 38], to estimate body state in real-time to control assistive devices [39, 40], and to predict effects of exoskeleton assistance [41] and surgical interventions [42] on muscle coordination. However, such simulations track recorded motions and do not produce new movement.

Alternatively, musculoskeletal simulations can produce motions without reference motion data using trajectory optimization methods. Instead of modeling the motor control system, this approach directly optimizes muscle activations that can actuate the musculoskeletal model and produce desired motions. While this approach does not take into account the structures or constraints of the human nervous system, the approach has been successful in reproducing well-practiced, or well-optimized, motor tasks. For example, normal walking and running motions can be produced by optimizing the muscle activations of a musculoskeletal model to move at a target speed with minimum muscle effort [43, 24, 25, 44]. Although this approach provides biomechanical insights by presenting theoretically optimal gaits for specific objectives, it is not suitable for predicting suboptimal behaviors that emerge from the underlying controller such as walking in unusual environments or reacting to unexpected perturbations.

#### Neuromechanical control models and simulations

A neuromechanical model includes a representation of a neural controller in addition to the musculoskeletal system (Fig. 1). To demonstrate that a controller can produce stable locomotion, neurome-chanical models are typically tested in a forward physics simulation for multiple steps while dynamically interacting with the environment (e.g., the ground and the gravitational force). Researchers have used neuromechanical simulations to test gait assistive devices before developing hardware [45, 46] and to understand how changes in musculoskeletal properties affect walking performance [26, 7]. Moreover, the control model can be directly used to control bipedal robots [47, 48, 49] and assistive devices [50, 45, 51].

Modeling human motor control is crucial for a predictive neuromechanical simulation. However, most of our current understanding of human locomotion control is extrapolated from experimental studies of simpler animals [52, 53] as it is difficult to measure and interpret the activity of the millions of neurons involved in human motor control. Therefore, human locomotion control models have been proposed based on a few structural and functional control hypotheses that are shared in many animals (Fig. 3). First, locomotion in many animals can be interpreted as a hierarchical structure with two layers, where the lower layer generates basic motor patterns and the higher layer sends commands to the lower layer to modulate the basic patterns [52]. It has been shown in some vertebrates, including cats and lampreys, that the neural circuitry of the spinal cord, disconnected from the brain, can produce steady locomotion and can be modulated by electrical stimulation to change speed, direction and gait [54, 55]. Second, the lower layer seems to consist of two control mechanisms: reflexes [56, 57] and central pattern generators (CPGs) [58, 59]. In engineering terms, reflexes and CPGs roughly correspond to feedback and feedforward control, respectively. Muscle synergy, where a single pathway co-activates multiple muscles, has also been proposed as a lower layer control mechanism that reduces the degrees of freedom for complex control tasks [60, 61]. Lastly, there is a consensus that humans use minimum effort to conduct well-practiced motor tasks, such as walking [62, 63]. This consensus provides a basis for using energy or fatigue optimization [24, 25, 26] as a principled means of finding control parameter values.

**Figure 3:**
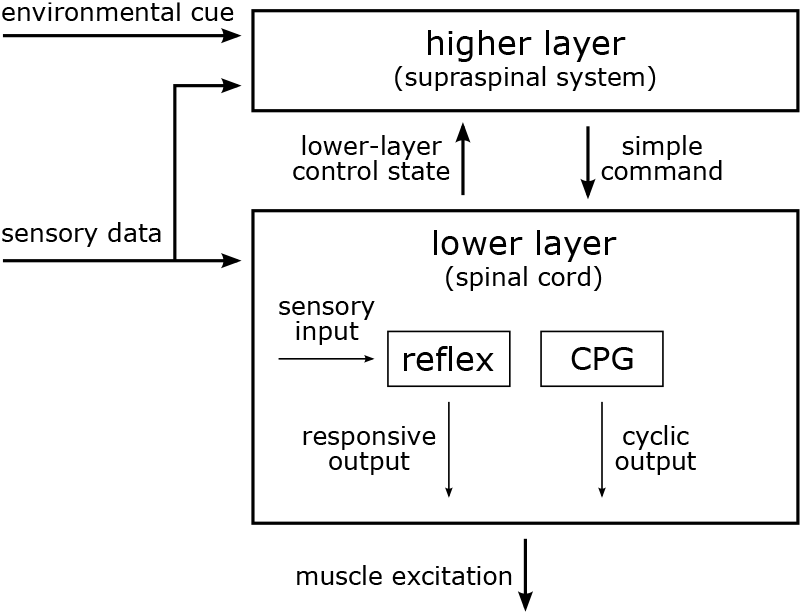
Locomotion control. The locomotion controller of animals is generally structured hierarchically with two layers. Reflexes and central pattern generators are the basic mechanisms of the lower layer controller.

Most neuromechanical control models are focused on lower layer control using spinal control mechanisms, such as CPGs and reflexes. CPG-based locomotion controllers consist of both CPGs and simple reflexes, where the CPGs, often modeled as mutually inhibiting neurons [64], generate the basic muscle excitation patterns. These CPG-based models [65, 66, 67, 68, 69, 8] demonstrated that stable locomotion can emerge from the entrainment between CPGs and the musculoskeletal system, which are linked by sensory feedback and joint actuation. CPG-based models also have been integrated with different control mechanisms, such as muscle synergies [68, 69, 8] and various sensory feedback circuits [66, 68]. On the other hand, reflex-based control models consist of simple feedback circuits without any temporal characteristics and demonstrate that CPGs are not necessary for producing stable locomotion. Reflex-based models [70, 18, 6, 71, 72] mostly use simple feedback laws based on sensory data accessible at the spinal cord such as proprioception (e.g., muscle length, speed and force) and cutaneous (e.g., foot contact and pressure) data [53, 57]. A reflex-based control model combined with a simple higher layer controller that regulates foot placement to maintain balance produced diverse locomotion behaviors including walking, running, and climbing stairs and slopes [6] and reacted to a range of unexpected perturbations similarly to humans [73] (Fig. 4). Reflex-based controllers also have been combined with CPGs [71] and a deep neural network that operates as a higher layer controller [72] for more control functions, such as speed and terrain adaptation.

**Figure 4:**
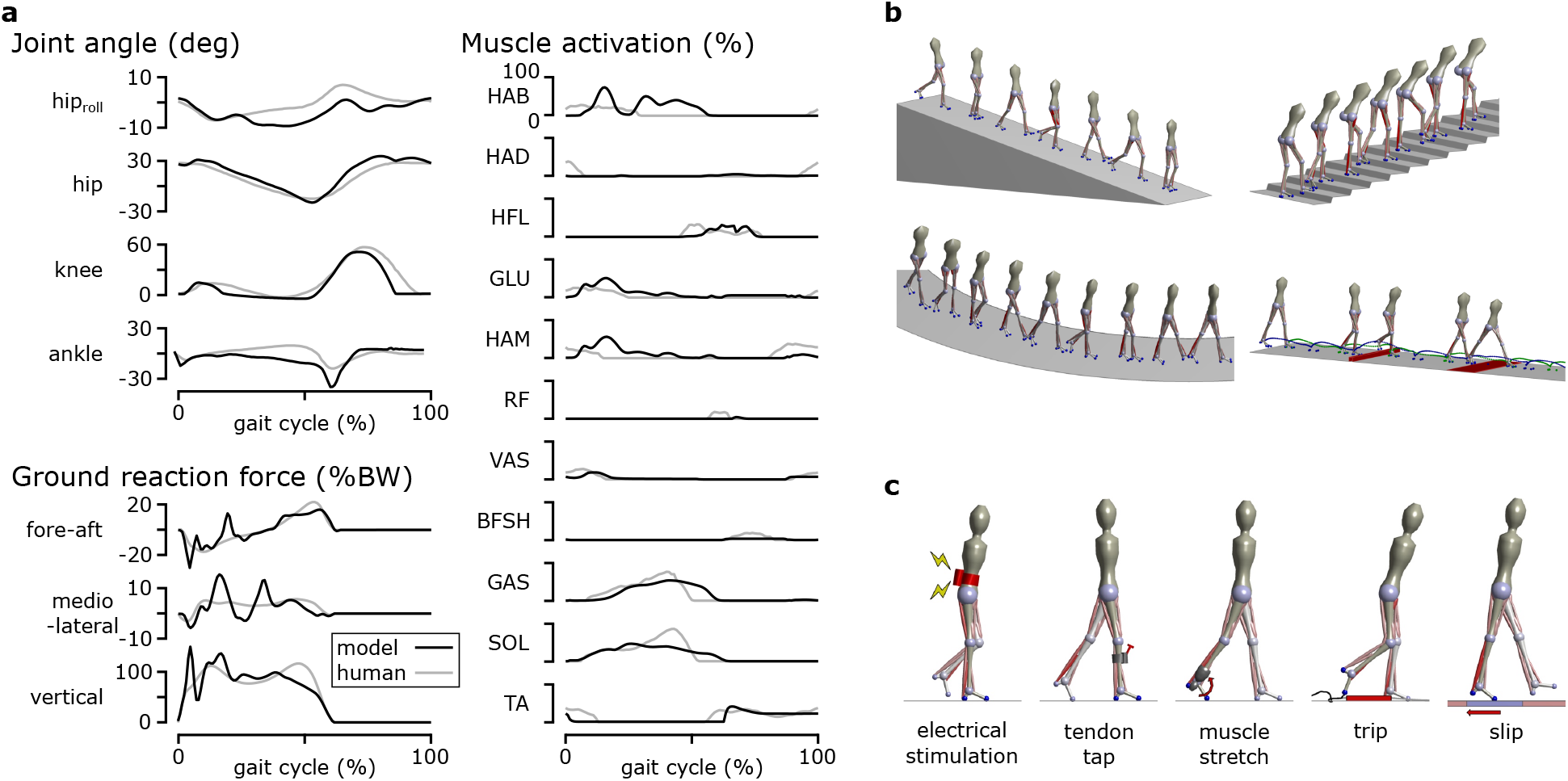
Reflex-based neuromechanical model. a. A reflex-based control model produced walking with human-like kinematics, dynamics, and muscle activations when optimized to walking with minimum metabolic energy consumption [6]. b. The model produced diverse locomotion behaviors when optimized at different simulation environments with different objectives. c. The same model optimized for minimum metabolic energy consumption reacted to various disturbances as observed in human experiments [73].

#### Human locomotion simulations for computer graphics

Human locomotion simulations have been widely developed in computer graphics to automate the process of generating human-like locomotion for computer characters. A number of controllers have been developed for characters to perform desired behaviors in physics simulations [74, 75, 76, 77, 78]. A variety of techniques have been proposed for simulating common behaviors, such as walking and running [79, 80, 81, 82]. Reference motions, such as motion capture data, were often used in the development process to produce more natural behaviors [83, 84, 85, 86]. Musculoskeletal models also have been used to achieve naturalistic motions [87, 88, 89], which makes them very close to neuromechanical simulations. The focus of these studies is producing natural motions rather than accurately representing the underlying biological system. However, the computer graphics studies and physiologically plausible neuromechanical simulations may converge as they progress to produce and model a wide variety of human motions.

#### Plausibility and limitations of control models

There are several ways to evaluate the physiological plausibility of control models. Plausibility of control models is of vital importance for the credibility of scientific knowledge the model produce and the implications of model predictions for rehabilitation research. Models that include neurophysiological details could be investigated more directly. For instance, a model that represents individual neurons and neural properties can be evaluated at the individual neuron level using neural recordings [90] and at the neuroanatomy level by assessing functions and connections of model parts that represent different areas of the brain and the spinal cord [91]. However, most of the control models tested in neuromechanical simulations focus on how motions emerge from motor control hypotheses rather than capturing the neural details. These models can be evaluated by the neuromechanical simulation results and the control features encoded in the models.

The plausibility of a neuromechanical control model can be assessed by the resulting simulation behavior. First of all, generating stable locomotion in neuromechanical simulations is a challenging control problem [92, 53] and thus has implications for the controller. For instance, a control model that cannot produce stable walking with physiological properties, such as nonlinear muscle dynamics and neural transmission delays, is likely missing some important aspects of human control [93]. Once motions are successfully simulated, they can be compared to measurable human data. We can say a model that produces walking with human-like kinematics, dynamics, and muscle activations is more plausible than one that does not. A model can be further compared with human control by evaluating its reactions to unexpected disturbances [73] and its adaptations in new conditions, such as musculoskeletal changes [26, 7], external assistance [45, 46], and different terrains.

We also can discuss whether a control model is constructed in a plausible manner. It is plausible for a control model to use sensory data that are known to be used in human locomotion [53, 57] and to work with known constraints, such as neural transmission delays. Models developed based on control hypotheses proposed by neuroscientists, such as CPGs and reflexes, partially inherit the plausibility of the corresponding hypotheses. Showing that human-like behaviors emerge from optimality principles that regulate human movements, such as minimum metabolic energy or muscle fatigue, also increases the plausibility of the control models [24, 25, 26].

Existing neuromechanical control models are mostly limited to modeling the lower layer control and producing steady locomotion behaviors. Most aspects of the motor learning process and the higher layer control are thus missing in current neuromechanical models. Motor learning occurs in daily life when acquiring new motor skills or adapting to environmental changes. For example, the locomotion control system adapts when walking on a slippery surface, moving a heavy load, walking on a split-belt treadmill [94, 95], and wearing an exoskeleton [96, 97]. The higher layer control processes environment cues, plans long-term motion strategies, and coordinates basic motor skills to navigate in dynamic and complex environments. While we will discuss other ideas for explicitly modeling motor learning and higher layer control in neuromechanical simulations in the sec:future section, deep RL may be an effective approach to developing controllers for challenging environments and motions.

### Deep reinforcement learning for motor control

This section highlights the concepts from deep reinforcement learning relevant to developing models for motor control. We provide a brief overview of the terminology and problem formulations of RL and then cover selected state-of-art deep RL algorithms that are relevant to successful solutions in the Learn to Move competition. We also review human locomotion simulation studies that have used deep RL.

#### Deep reinforcement learning

Reinforcement learning is a machine learning paradigm for solving decision-making problems. The objective is to learn an optimal policy *π* that enables an agent to maximize its cumulative reward through interactions with its environment [98] (Fig. 5). For example, in the case of the Learn to Move competition, the environment was the musculoskeletal model and physics-based simulation environment, and higher cumulative rewards were given to solutions that better followed target velocities with lower muscle effort. Participants developed agents, which consists of a policy that controls the musculoskeletal model and a learning algorithms that trains the policy. For the general RL problem, at each timestep *t*, the agent receives an observation *o_t_* (perception and proprioception data in the case of our competition; perception data includes information on the target velocities) and queries its policy *π* for an action *a_t_* (excitation values of the muscles in the model) in response to that observation. An observation *o_t_* is the full or partial information of the state *s_t_* of the environment. The policy *π*(*a_t_|o_t_*) can be either deterministic or stochastic, where a stochastic policy defines a distribution over actions *a_t_* given a particular observation *o_t_* [99]. The agent then applies the action in the environment, resulting in a transition to a new state *s_t_*_+1_ and a scalar reward *r_t_* = *r*(*s_t_, a_t_, s_t_*_+1_). The state transition is determined according to the dynamics model *ρ*(*s_t_*_+1_ *s_t_, a_t_*). The objective for the agent is to learn an optimal policy that maximizes its cumulative reward, or return.

**Figure 5:**
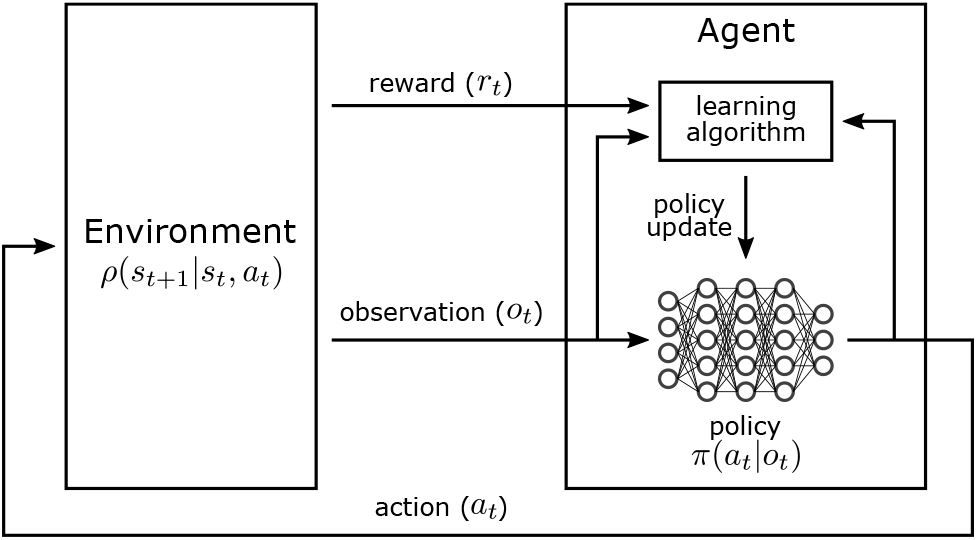
Reinforcement learning. In a typical RL process, an agent takes a reward and observation as input and trains a policy that outputs an action to achieve high cumulative rewards.

One of the crucial design decisions in applying RL to a particular problem is the choice of policy representation. While a policy can be modeled by any class of functions that maps observations to actions, the recent use of deep neural networks to model policies demonstrated promising results in complex problems and has led to the emergence of the field of deep RL. Policies trained with deep RL methods achieved human-level performance on many of the 2600 Atari video games [100] and overtook world champion human players in the game of Go [101, 102].

#### State-of-the-art deep RL algorithms used in Learn to Move

Model-free deep RL algorithms (Fig. 6) are widely used for continuous control tasks, such as those considered in the Learn to Move competition where the actions are continuous values of muscle excitations. Model-free algorithms do not learn an explicit dynamics model of state transitions; instead, they directly learn a policy to maximize the expected return. In these continuous control tasks, the policy specifies actions that represent continuous quantities such as control forces or muscle excitations. Policy gradient algorithms incrementally improve a policy by first estimating the gradient of the expected return using trajectories collected from rollouts (forward simulations in our case) of the policy, and then updating the policy via gradient ascent [103]. While simple, the standard policy gradient update has several drawbacks, including stability and sample efficiency. First, the gradient estimator can have high variance, which can lead to unstable learning, and a good gradient estimate may require a large number of training samples. Popular algorithms such as TRPO [104] and PPO [105] improve the stability of policy gradient methods by limiting the change in the policy’s behavior after each update step, as measured by the relative entropy between the policies [106]. Another limitation of policy gradient methods is their low sample efficiency, often requiring millions of samples to solve relatively simple tasks. Off-policy gradient algorithms, which utilize rollouts collected from previous policies, have been proposed to substantially reduce the number of samples required to learn effective policies [107, 108, 109]. Off-policy algorithms, such as DDPG [107], typically fit a Q-function, *Q*(*s, a*), which is the expected return of performing an action *a* in the current state *s*. These methods differentiate the learned Q-function to approximate the policy gradient, then use it to update the policy. Recent off-policy methods, such as TD3 and SAC, build on this approach and propose several modifications that further improve sample efficiency and stability.

**Figure 6:**
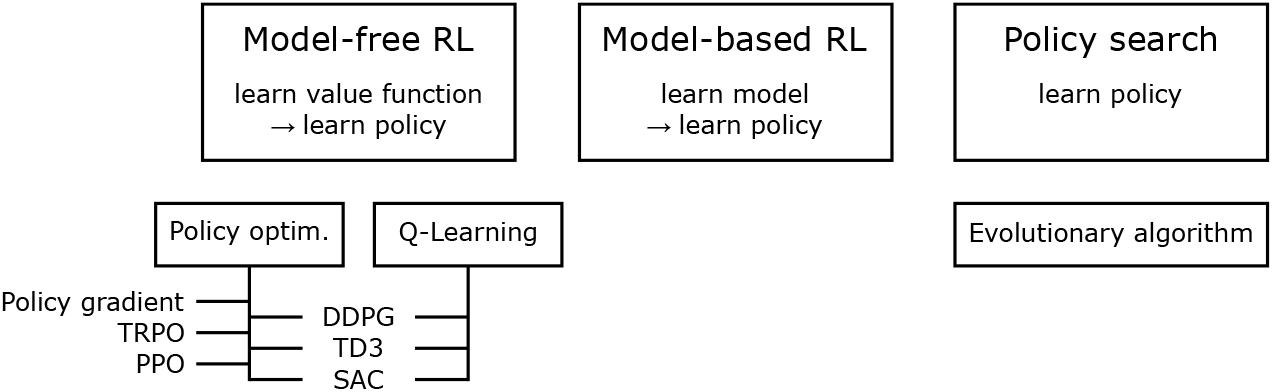
Reinforcement learning algorithms for continuous action space. The diagram is adapted from [110] and presents a partial taxonomy of RL algorithms for continuous control, or continuous action space. This focuses on a few modern deep RL algorithms and some traditional RL algorithms that are relevant to the algorithms used by the top teams in our competition. TRPO: trust region policy optimization [104]; PPO: proximal policy optimization [105]; DDPG: deep deterministic policy gradients [107]; TD3: twin delayed deep deterministic policy gradients [108]; SAC: soft-actor critic [109].

#### Deep RL for human locomotion control

Human motion simulation studies have used various forms of RL (Fig. 6). A number of works in neuromechanical simulation [67, 6] and computer graphics studies [87, 88] reviewed in the sec:simLoco section used policy search methods [111] with derivative-free optimization techniques, such as evolutionary algorithms, to tune their controllers. The control parameters are optimized by repeatedly running a simulation trial with a set of control parameters, evaluating the objective function from the simulation result, and updating the control parameters using an evolutionary algorithm [112]. This approach makes very minimal assumptions about the underlying system and can be effective for tuning controllers to perform a diverse array of skills [113, 6]. However, these algorithms often struggle with high dimensional parameter spaces (i.e., more than a couple of hundred parameters), therefore some care is required to design controllers that expose a relatively low-dimensional but expressive set of parameters for optimization. Selecting an effective set of parameters can require a great deal of expertise, and the selected set of parameters tend to be specific for particular skills, limiting the behaviors that can be reproduced by the character.

Recently, deep RL techniques have demonstrated promising results for character animation, with policy optimization methods emerging as the algorithms of choice for many of these applications [103, 107, 105]. These methods have been effective for training controllers that can perform a rich repertoire of skills [114, 10, 115, 116, 117, 118]. One of the advantages of deep RL techniques is the ability to learn controllers that operate directly on high-dimensional, low-level representations of the underlying system, thereby reducing the need to manually design compact control representations for each skill. These methods have also been able to train controllers for interacting with complex environments [114, 119, 120], as well as for controlling complex musculoskeletal models [121, 11]. Reference motions continue to play a vital role in producing more naturalistic behavior in deep RL as a form of deep imitation learning, where the objective is designed to train a policy that mimics human motion capture data [10, 117, 11] (Fig. 7). These studies show the potential of using deep RL methods to develop versatile controllers for musculoskeletal models and studying human motor control.

**Figure 7:**
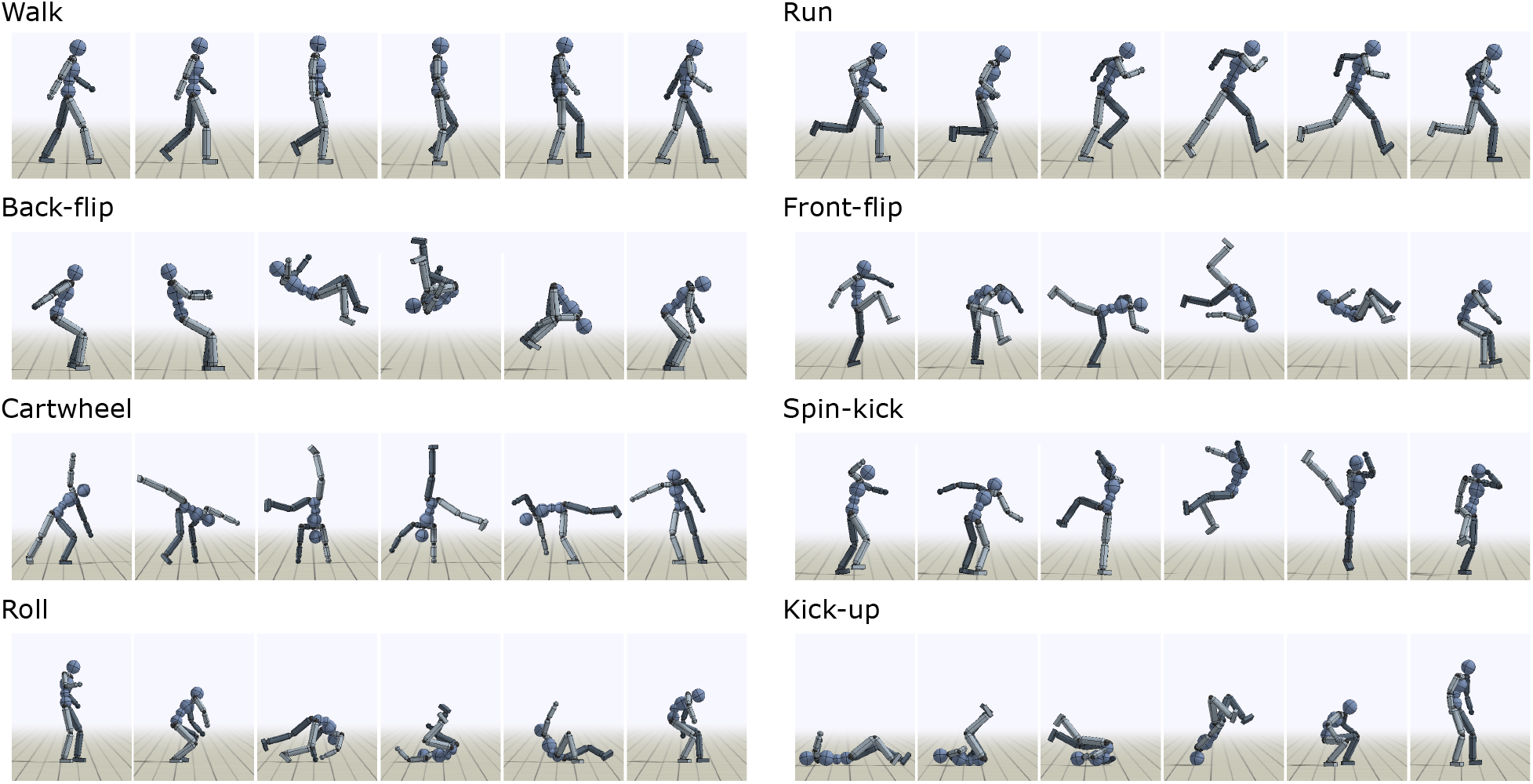
Computer graphics characters performing diverse human motions. Dynamic and acrobatic skills learned to mimic motion capture clips with RL in physics simulation [10].

### Learn to Move competition

The potential synergy of neuromechanical simulations and deep RL methods in modeling human control motivated us to develop the OpenSim-RL simulation platform and to organize the Learn to Move competition series. OpenSim-RL [33] leverages OpenSim to simulate musculoskeletal models and OpenAI Gym, a widely used RL toolkit [122], to standardize the interface with state-of-the-art RL algorithms. OpenSim-RL is open-source and is provided as a Conda package [123], which has been downloaded about 42,000 times (counting multiple downloads from the same users) over the past three years. Training a controller for a human musculoskeletal model is a difficult RL problem considering the large-dimensional observation and action spaces, delayed and sparse rewards resulting from the highly non-linear and discontinuous dynamics, and the slow simulation of muscle dynamics. Therefore, we organized the Learn to Move competition series to crowd-source machine learning expertise in developing control models of human locomotion. The mission of the competition series is to bridge neuroscience, biomechanics, robotics, and machine learning to model human motor control.

The Learn to Move competition series has been held annually since 2017. It has been one of the official competitions at the NeurIPS conference, a major event at the intersection of machine learning and computational neuroscience. The first competition was NIPS 2017: Learning to Run [33, 124], and the task was to develop a controller for a given 2D human musculoskeletal model to run as fast as possible while avoiding obstacles. In the second competition, NeurIPS 2018: AI for Prosthetics Challenge [125], we provided a 3D human musculoskeletal model, where one leg was amputated and replaced with a passive ankle-foot prosthesis. The task was to develop a walking controller that could follow velocity commands, the magnitude and direction of which varied moderately. These two competitions together attracted about 1000 teams, primarily from the machine learning community, and established successful RL techniques which will be discussed in the sssec:solutions section. We designed the 2019 competition to build on knowledge gained from past competitions. For example, the challenge in 2018 demonstrated the difficulty of moving from 2D to 3D. Thus, to focus on controlling maneuvering in 3D, we designed the target velocity to be more challenging, while we removed the added challenge of simulating movement with a prosthesis. We also refined the reward function to encourage more natural human behaviors.

#### NeurIPS 2019: Learn to Move - Walk Around Overview

NeurIPS 2019: Learn to Move - Walk Around was held online from June 6 to November 29 in 2019. The task was to develop a locomotion controller, which was scored based on its ability to meet target velocity vectors when applied in the provided OpenSim-RL simulation environment. The environment repository was shared on Github [126], the submission and grading were managed using the AIcrowd platform [127], and the project homepage provided documentation on the environment and the competition [128]. Participants were free to develop any type of controller that worked in the environment. We encouraged approaches other than brute force deep RL by providing human gait data sets of walking and running [129, 130, 131] and a 2D walking controller adapted from a reflex-based control model [6] that could be used for imitation learning or in developing a hierarchical control structure. There were two rounds. The top 50 teams in Round 1 were qualified to proceed to Round 2 and to participate in a paper submission track. RL experts were invited to review the papers based on the novelty of the approaches, and we selected the best and finalist papers based on the reviews. The prizes included GPUs (NVIDIA), a motion capture suit (Xsens), travel grants, and an invitation to submit a paper to the Journal of NeuroEngineering and Rehabilitation and the NeurIPS 2019 Deep RL Workshop.

In total, 323 teams participated in the competition and submitted 1448 solutions. In Round 2, the top three teams succeeded in completing the task and received high scores (mean total rewards larger than 1300 out of 1500). Five papers were submitted, and we selected the best paper [132] along with two more finalist papers [133, 134]. The three finalist papers came from the top three teams, where the best paper was from the top team.

#### Simulation environment

The OpenSim-RL environment included a physics simulation of a 3D human musculoskeletal model, target velocity commands, a reward system, and a visualization of the simulation (Fig. 8). The 3D musculoskeletal model had seven segments connected with eight rotational joints and actuated by 22 muscles. Each foot segment had three contact spheres that dynamically interacted with the ground. A user-developed policy could observe 97-dimensional body sensory data and 242-dimensional target velocity map and produced a 22-dimensional action containing the muscle excitation signals. The reward was designed to give high total rewards for solutions that followed target velocities with minimum muscle effort. The details of the reward were set to result in human-like walking based on previous neuromechanical simulation studies [6, 26]. The mean total reward of five trials with different target velocities was used for ranking. The full description of the simulation environment is provided on our project homepage [135].

**Figure 8:**
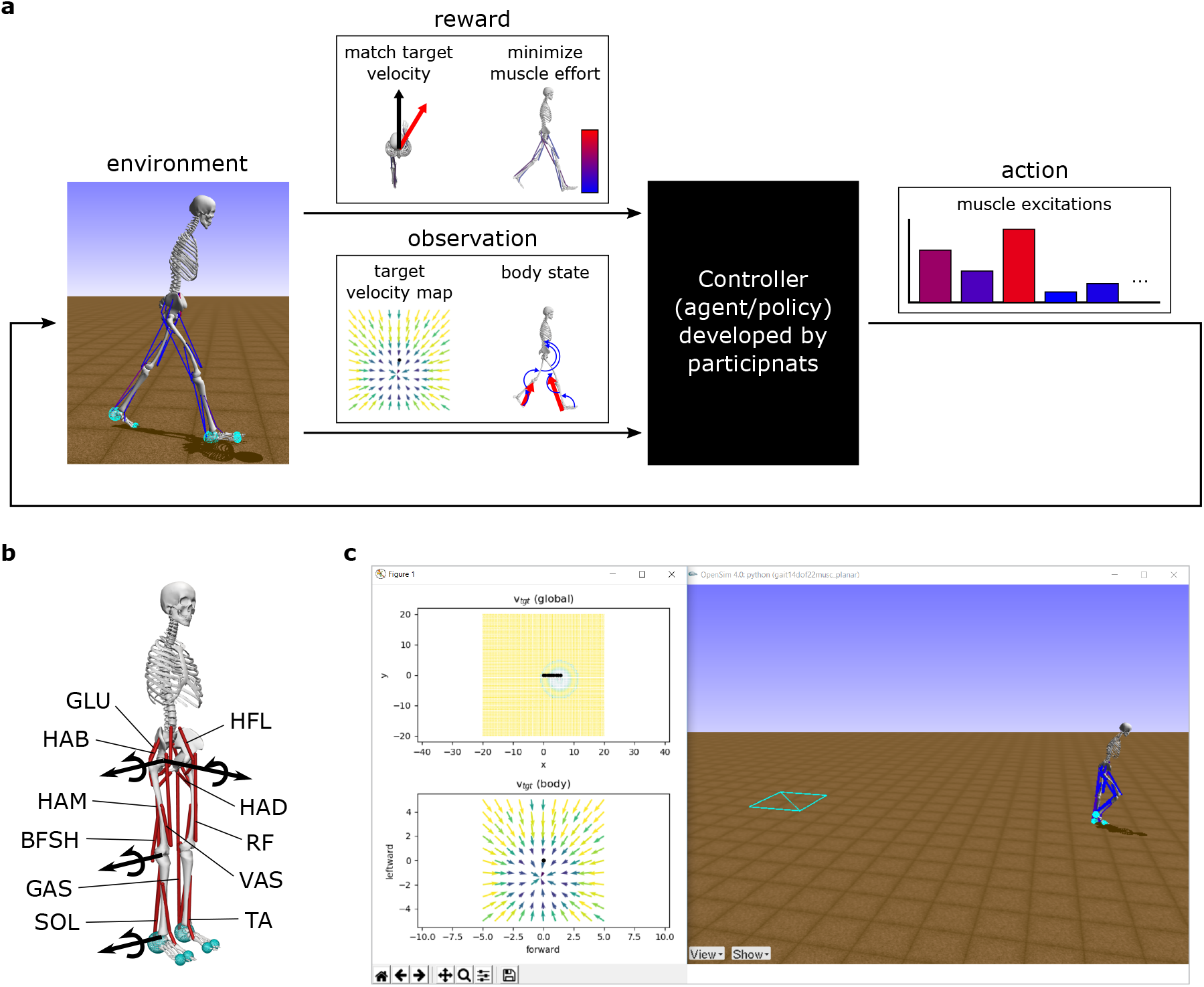
OpenSim-RL environment for the NeurIPS 2019: Learn to Move – Walk Around competition. a. A neuromechanical simulation environment is designed for a typical RL framework (Fig. 5). The environment took an action as input, simulated a musculoskeletal model for one time-step, and provided the resulting reward and observation. The action was excitation signals for the 22 muscles. The reward was designed so that solutions following target velocities with minimum muscle effort would achieve high total rewards. The observation consisted of a target velocity map and information on the body state. b. The environment included a musculoskeletal model that represents the human body. Each leg consisted of four rotational joints and 11 muscles. (HAB: hip abductor; HAD: hip adductor; HFL: hip flexor; GLU: glutei, hip extensor; HAM: hamstring, biarticular hip extensor and knee flexor; RF: rectus femoris, biarticular hip flexor and knee extensor; VAS: vastii, knee extensor; BFSH: short head of biceps femoris, knee flexor; GAS: gastrocnemius, biarticular knee flexor and ankle extensor; SOL: soleus, ankle extensor; TA: tibialis anterior, ankle flexor) c. The simulation environment provided a real-time visualization of the simulation to users. The global map of target velocities is shown at the top-left. The bottom-left shows its local map, which is part of the input to the controller. The right visualizes the motion of the musculoskeletal model.

#### Top solutions and results

Various RL techniques have been effectively used since the first competition [124, 125], including frame skipping, discretization of the action space, and reward shaping. These are practical techniques that constrain the problem in certain ways to encourage an agent to search successful regions faster in the initial stages of training. Frame skipping repeats a selected action for a given number of frames instead of operating the controller every frame [133]. This technique reduces the sampling rate and thus computations while maintaining a meaningful representation of observations and control. Discretization of the muscle excitations constrains the action space and thus the search space, which can lead to much faster training. In the extreme case, binary discretization (i.e., muscles were either off or fully activated) was used by some teams in an early stage of training. Reward shaping modifies the reward function provided by the environment to encourage an agent to explore certain regions of the solution space. For example, a term added to the reward function that penalizes crossover steps encouraged controllers to produce more natural steps [133, 134]. Once agents found solutions that seem to achieve intended behaviors with these techniques, they typically were further tuned with the original problem formulation.

A common trend across the finalists in 2019 was the use of off-policy methods. The first place entry by Zhou et al. [132] used DDPG [107], the second place entry by Kolesnikov and Hrinchuk [133] used TD3 [108], and the third place entry by Akimov [134] used SAC [109]. Since off-policy algorithms use data collected with previous policies for training, they can be substantially more sample efficient than their on-policy counterparts and could help to compensate for the computationally expensive simulation. Off-policy algorithms are also more amenable to distributed training, since data-collection and model updates can be performed asynchronously. Kolesnikov and Hrinchuk [133] leveraged this property of off-policy methods to implement a population-based distributed training framework, which used an ensemble of agents whose experiences were collected into a shared replay buffer that stored previously collected *(observation, action, reward, next observation)* pairs. Each agent was configured with different hyperparameter settings and was trained using the data collected from all agents. This, in turn, improved the diversity of the data that was used to train each policy and also improved the exploration of different strategies for solving a particular task.

Curriculum learning [136] was also used by the top teams. Curriculum learning is a training method where a human developer designs a curriculum that consists of a series of simpler tasks that eventually lead to the original task that is challenging to train from scratch. Akimov [134] first trained a policy to walk and then to follow target velocities. Similarly, Zhou et al. [132] trained a policy for normal speed walking by first training it to run at high speed, then to run at slower speeds, and eventually to walk at normal speed. They found that the policy trained through this process resulted in more natural gaits than policies that were directly trained to walk at normal speeds. This is probably because there is a limited set of very high-speed gaits that are close to human sprinting, and starting from this human-like sprinting gait could have guided the solution to a more natural walking gait out of a large variety of slow gaits, some of which are unnatural and ineffective local minima. Then they obtained their final solution policy by training this basic walking policy to follow target velocities and to move with minimum muscle effort.

The winning team, Zhou et al., proposed risk averse value expansion (RAVE), a hybrid approach of model-based and model-free RL [132]. Their method fits an ensemble of dynamics models (i.e. models of the environment) to data collected from the agent’s interaction with the environment, and then uses the learned models to generate imaginary trajectories for training a Q-function. This model-based approach can substantially improve sample efficiency by synthetically generating a large volume of data but can also be susceptible to bias from the learned models, which can negatively impact performance. To mitigate potential issues due to model bias, RAVE uses an ensemble of dynamics models to estimate the confidence bound of the predicted values and then trains a policy using DDPG to maximize the confidence lower bound. Their method achieved impressive results on the competition tasks and also demonstrated competitive performance on standard OpenAI Gym benchmarks [122] compared to state-of-the-art algorithms [132].

#### Implications for human locomotion control

The top solution [132] shows that it is possible to produce many locomotion behaviors with the given 3D human musculoskeletal model, despite its simplifications. The musculoskeletal model simplifies the human body by, for example, representing the upper body and the pelvis as a single segment. Moreover, the whole body does not have any degree of freedom for internal yaw motion (Fig. 8-a). Such a model was selected for the competition as it can produce many locomotion behaviors including walking, running, stair and slope climbing, and moderate turning as shown in a previous study [6]. On the other hand, the missing details of the musculoskeletal model could have been crucial for generating other behaviors like sharp turning motions and gait initiation. However, the top solution was able to initiate walking from standing, quickly turn towards a target (e.g., turn 180^◦^ in one step; Fig. 9), walk to the target at commanded speeds, and stop and stand at the target. To our knowledge, it is the first demonstration of rapid turning motions with a musculoskeletal model with no internal yaw degree of freedom. The solution used a strategy that is close to a step-turn rather than a spin-turn, and it will be interesting to further investigate how the simulated motion compares with human turning [137, 138].

**Figure 9:**
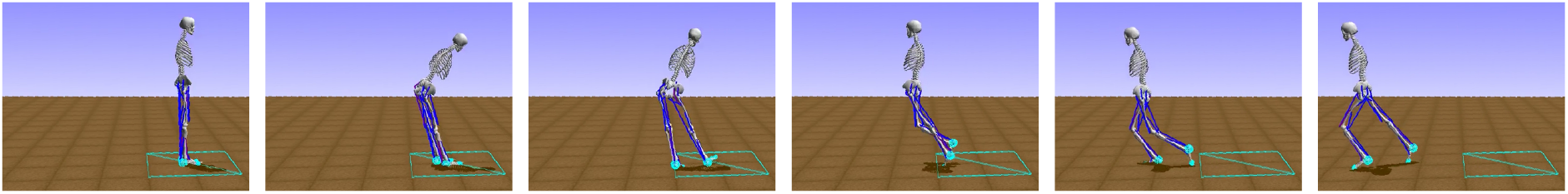
Rapid turning motion. The top solution can make the musculoskeletal model with no internal yaw degree of freedom to turn 180° in a single step. Snapshots were taking every 0.4 s.

The top solutions had some limitations in producing human-like motions. In the top solution [132], the human model first turned to face the target then walked forward towards the target with a relatively natural gait. However, the gait was not as close to human walking as motions produced by previous neuromechanical models and trajectory optimization [6, 44]. This is not surprising as the controllers for the competition needed to cover a broad range of motions, and thus were more difficult to fully optimize for specific motions. The second and third solutions [133, 134] were further from human motions as they gradually moved towards a target often using side steps. As policy gradient methods use gradient descent, they often get stuck at local optima resulting in suboptimal motions [120] even though natural gaits are more efficient and agile. Although the top solution overcame some of these suboptimal gaits through curriculum learning, better controllers could be trained by utilizing imitation learning for a set of optimal motions [10, 11, 12] or by leveraging control models that produce natural gaits [18, 6]. Different walking gaits, some of which are possibly suboptimal, are also observed in toddlers during the few months of extensive walking experience [139, 140], and interpreting this process with an RL framework will be instructive to understanding human motor learning.

### Future directions

Various deep reinforcement learning approaches, such as imitation learning and hierarchical learning, approaches could be used to produce more complex motions and their coordination. The top solutions in the competitions were able to control standing, walking and running behaviors, which is impressive but only covers a small portion of human motion. Parkour athletes, for example, can plan and execute jumping, vaulting, climbing, rolling, and many other acrobatic motions to move in complex environments, which would be very difficult to perform with brute force RL methods. Imitation learning [10, 117, 11] could be used to train multiple networks to master a set of acrobatic skills (Fig. 7). These networks of motion primitives can then be part of a hierarchical controller [141, 142, 143], where a higher layer controller coordinates the motion skills. More control layers that analyze dynamic scenes and plan longer-term motion sequences [144, 145] can be added if a complex strategy is required for the task. We will promote research in developing all these aspects of control by designing a contest that requires complex motions and long-term motion planning. The task can be something like the World Chase Tag competition, where two athletes take turns to tag the opponent, using athletic movements, in an arena filled with obstacles [146]. To this end, we plan to develop a simulation environment with a faster physics engine that can handle multi-body contacts between human models and obstacles [3, 4, 5], provide a human musculoskeletal model with an articulated upper body [88, 11], and design a visuomotor control interface [147].

As discussed in the ssec:plausibility section, the physiologically plausible control models are currently limited to representing the lower layer control and omit many aspects of the motor learning process and the higher layer control. Motor learning paradigms have been proposed to explain the transient behaviors observed while acquiring new motor skills or adapting to environment changes [148, 149]. Integrating these paradigms into locomotion control models will substantially expand the realm of human motions that neuromechanical simulations can represent. Gait adaptation reported during walking on a split-belt treadmill [94, 95] are a benchmark for evaluating motor learning simulations, and simulations that correctly predict the training process of walking with exoskeletons [96, 97] could change the way we develop assistive devices. Deep RL methods may be able to train control models that perform the functions of both higher layer and lower layer controls. However, integrating physiologically plausible control models with deep RL methods can have both predictive and computational advantages. As there are CPG- and reflex-based lower layer control models proposed to represent some of the physiological control constraints [67, 6], combining these lower-layer control models with a high-layer controller developed with deep RL may have some utility in producing motions closer to humans in novel scenarios. Moreover, it may be easier to train this higher-layer deep neural network than one without such structured lower-layer control as the solution space could be constrained to reasonable movements. Similarly, the action space can be reduced by having a muscle synergy layer between the deep neural network and muscle excitations. In future competitions, we plan to implement and share more physiologically plausible control models in OpenSim-RL and organize a separate paper submission track to acknowledge approaches that encode physiological features.

The importance of physiological plausibility of predictive simulations is not limited to control models and simulation results but also applies to the musculoskeletal simulation environment. For instance, there are a variety of muscle models [20, 150, 151] that represent biological muscles at different levels of detail and focus on different properties, and investigating the effect of these models in whole-body motion will be instructive. Body motion can be affected by other physical and physiological details as well, such as the number of muscles and body segments used to represent the musculoskeletal system [11], soft tissue [152] and its interaction with assistive devices [153], muscle fatigue [154], and musculoskeletal properties that vary across individuals [155] and health conditions [156, 157]. Modeling these details requires sufficient domain knowledge in each field. In the long term, we aim to modularize our simulation framework so that it will be easier for researchers to focus on their expertise, for example in implementing and testing different muscle models, while using other modules (e.g., control, training, and environment modules) in our framework. We could collect various control models and RL algorithms through the Learn to Move competition and share them as part of the modular framework that will make motion simulations easier for non-RL experts.

## Conclusion

In this article, we reviewed neuromechanical simulations and deep reinforcement learning with a focus on human locomotion. Neuromechanical simulations provide a means to evaluate control models, and deep RL is a promising tool to develop control models for complex movements. We hope to see more interdisciplinary studies and collaboration that uses advanced machine learning techniques to develop and evaluate physiologically plausible control models. Such studies could significantly advance our ability to model control and predict human behaviors. We plan to continue to develop and disseminate the Learn to Move competition and its accompanying simulation platform to facilitate these advancements toward predictive neuromechanical simulations for rehabilitation treatment and assistive devices.

## Abbreviations

CPG: central pattern generators
RAVE: risk averse value expansion
RL: reinforcement learning

## Acknowledgements

We thank our partners and sponsors: AIcrowd, Google, Nvidia, Xsens, Toyota Research Institute, the NeurIPS Deep RL Workshop team, and the Journal of NeuroEngineering and Rehabilitation. We thank all the external reviewers who provide us valuable feedback on the papers submitted to the competition. And most importantly, we thank all the researchers who have competed in the Learn to Move competition.

## Author’s contributions

All authors are members of the team that organized the Learn to Move competition. SS, LK, XBP, CO and JH drafted the manuscript. All authors edited and approved the submitted manuscript.

## Funding

The Learn to Move competition series were co-organized by the Mobilize Center, a National Institutes of Health Big Data to Knowledge Center of Excellence supported through Grant U54EB020405. The first author was supported by the National Institutes of Health under Grant K99AG065524.

## Availability of data and materials

The environment repository and documentation of NeurIPS 2019: Learn to Move - Walk Around is shared on Github [126] and the project homepage [128].

## Ethics approval and consent to participate

Not applicable.

## Consent for publication

Not applicable.

## Competing interests

The authors declare that they have no competing interests.

## Notes

### Competing Interest Statement

The authors have declared no competing interest.

